# CARTiBASE: an interactive knowledge base for CAR sequence retrieval and similarity analysis

**DOI:** 10.64898/2026.02.25.707638

**Authors:** Gauthier Le Compte, Halil Ceylan, Pieter Meysman, Kris Laukens

**Affiliations:** Adrem Data Lab, Department of Computer Science, University of Antwerp, Antwerp, Belgium; Antwerp Unit for Data Analysis and Computation in Immunology and Sequencing (AUDACIS), University of Antwerp, Antwerp, Belgium; Biomedical Informatics Research Network Antwerp (Biomina), University of Antwerp, Antwerp, Belgium

**Author notes:** **Corresponding Author:** Kris Laukens. Shared senior authors (Pieter Meysman, Kris Laukens).

## Abstract

**Summary:** Chimeric Antigen Receptors (CARs) are modular synthetic constructs that have transformed cellular immunotherapy, enabling targeted recognition and killing of malignant cells. Their clinical success has driven an explosive growth in new receptor designs, but these sequences are dispersed across heterogeneous sources such as publications, patents and supplementary files. This fragmentation and inconsistency limits comparative analysis, reproducibility and the reuse of existing constructs. To address this, we curated and standardized more than 10,000 CAR sequences into a single, harmonized resource. CARTiBASE is a web-based platform that provides standardized annotation, interactive browsing and fast similarity search across this curated collection. This unique database was leveraged to analyse the diversity in current CAR constructs within the public domain, revealing common design trends and lineages, as well as highlighting potential avenues for future CAR development.

**Availability and Implementation:** CARTiBASE is freely available for non-commercial use at https://www.cartibase.org, without mandatory registration. The web server is implemented with a Python/Flask API backend and a Vue-based frontend and supports all major browsers.

Users can search and filter thousands of CARs, inspect domain boundaries across signal peptide, antigen-binding domain, hinge, transmembrane, co-stimulatory and intracellular signaling regions, compare constructs and download sequences as FASTA files for downstream use.

## Introduction

CAR-T cell therapy is a form of adoptive immunotherapy in which patient or donor T cells are engineered to express a **Chimeric Antigen Receptor (CAR)** that enables antigen-specific recognition of malignant cells [1]. A CAR is constructed from several modular components that together determine antigen specificity, structural configuration and signaling output. These typically include: (i) an extracellular **antigen-binding domain**, which allows recognition of a target antigen expressed on the tumor cells surface, (ii) a **hinge domain** that provides flexibility and optimal positioning for antigen binding, (iii) a **transmembrane domain** which anchors the CAR within the T cell membrane and influences receptor stability, and (iv) several **intracellular signaling domains** that dictate the strength, persistence and functionality of the CAR-T cell response [2].

The last decade has seen a rapid expansion in CAR engineering strategies, resulting in substantial **diversity across all domains** of the receptor. Variants differ in scFv origin, hinge length and composition, choice of transmembrane region, and combinations of co-stimulatory and signaling domains. This **heterogeneity** complicates comparative analysis, as domain choices strongly influence CAR expression, tonic signaling, antigen sensitivity and therapeutic performance [3, 4]. Despite the growing number of CAR constructs, their sequences are not stored in a centralized resource and are frequently reported in **inconsistent formats** such as **partial sequences, mixed annotations** or **non-machine-readable files**. This fragmentation hinders reproducibility, complicates comparative analyses and limits systematic exploration of how CAR designs vary across domains. To address these challenges, we developed **CARTiBASE**, an interactive database and online platform that unifies CAR sequences into a **standardized representation** and provides **domain-level annotation and similarity search**. CARTiBASE enables comparative analysis of CAR designs through a consistent and accessible interface and supports reuse of existing CAR constructs for both computational and experimental studies.

## Implementation

### Data model and storage

CARTiBASE stores each Chimeric Antigen Receptor as a **standardized record**. Internally, all CARs are maintained in a curated CSV file in which each row corresponds to a single construct, and all columns capture (i) the full amino acid sequence, (ii) the segmented domain sequences and boundaries, (iii) target antigen information when available, and (iv) source metadata, including contributing authors and affiliated institutions. This flat-file format allows for **fast loading, simple filtering** and **full transparency** of the underlying data. All records can be viewed through the web interface and downloaded as FASTA files for offline or computational analysis.

### Domain segmentation and target antigen annotation

CARTiBASE segments each CAR into four sequential components: the signal peptide, the extracellular region, the transmembrane domain and the intracellular region. Segmentation is performed via a stepwise annotation strategy that integrates **DeepTMHMM** and **HMMER-based profile** searches [5, 6]. In parallel, target antigen information is extracted where available using a lightweight rule-guided workflow that identifies antigen names from titles, publication headers and additional notes (*Figure 1*).

**Figure 1.**
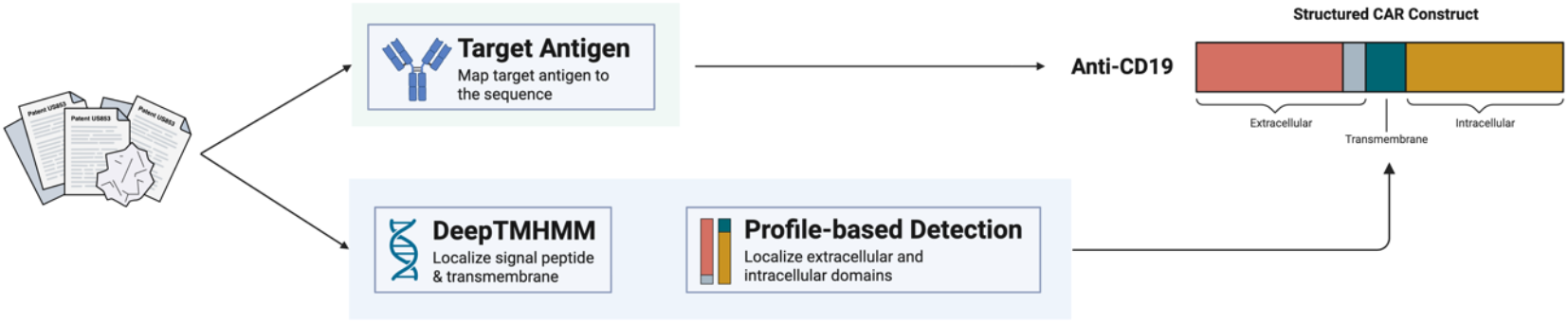
Domain segmentation and target antigen annotation workflow. Created in BioRender.

First, **DeepTMHMM**, a deep learning–based transmembrane topology predictor, is used to identify signal peptides, membrane-spanning helices and cytosolic regions. The model outputs a per-residue topology label and associated probabilities, allowing for the precise localization of (i) the signal peptide, (ii) the boundary between extracellular and transmembrane regions and (iii) the start of the intracellular tail. Next, the extracellular and intracellular regions are analyzed using **HMMER** against curated sets of profile Hidden Markov Models (HMMs). These profiles include models for common extracellular antigen-binding modules (e.g. scFv-related VH/VL motifs) and common intracellular signaling domains (e.g. CD28, 4-1BB, CD3ζ). HMMER returns domain hits with alignment coordinates, bit scores and E-values, and only high-confidence matches are retained and mapped onto the DeepTMHMM-derived topology.

**Target antigen annotation** is performed using **a rule-based pattern matching pipeline** that combines **keyword detection** (e.g. “anti-…”, “CD\d+”) with **regular expressions** and a curated dictionary of known antigens. Candidate terms extracted from text metadata are normalized and mapped to a pre-defined vocabulary, producing a standardized antigen field for each CAR construct. For sequences lacking explicit antigen information, CARTiBASE currently leaves this field unassigned; however, future versions will incorporate sequence-based inference strategies.

### Similarity search and alignment details

CARTiBASE includes a fast similarity search module based on a **DIAMOND-compatible workflow** [7]. All CAR protein sequences from the master file are compiled into a precomputed DIAMOND database that can be queried with either full-length CARs or individual domain sequences. Query sequences must be at least 15 amino acids long for the similarity search to be performed. For a given query sequence or uploaded FASTA file, the server runs DIAMOND against this database and returns ranked hits with identity, coverage and E-values, together with alignment coordinates. The web interface displays the results as sortable hit tables with links to the corresponding CARTiBASE entries and alignment details for sequence comparison.

### Web application and user interface

The CARTiBASE web application consists of a **Python/Flask** backend and a frontend implemented in **Vue**. The backend handles loading and querying of the CSV data file, retrieval of CAR records and domain annotations, execution of similarity searches and generation of FASTA downloads. The Vue frontend offers (i) global search and filtering, (ii) sequence pages with domain maps and annotations, (iii) similarity search with alignment details and (iv) FASTA downloads. Users can search by accession number, patent title or amino acid sequence, or alternatively upload a FASTA file and select one sequence to use as the search query.

## Results

### Database content and coverage

CARTiBASE collects CAR sequences from public sources and standardizes them into a consistent representation for downstream analysis. Each entry includes the full amino acid sequence, domain segmentation, target information when available and the associated publication or patent reference. The database is continuously updated as new CAR designs appear in the literature and patent repositories. The database is updated on a monthly basis to incorporate newly published or patented CAR constructs. In its first deployment, CARTiBASE contains 10,979 CAR sequences targeting a broad spectrum of antigens and featuring a wide range of domain combinations. BCMA and CD19 dominate the dataset, followed by CD22, CD123 and CD20, together forming the top five most frequently targeted antigens (Figure 2). These target frequencies reflect the annotations derived by our rule-based extraction pipeline from the available metadata and do not rely on sequence-based inference.

**Figure 2.**
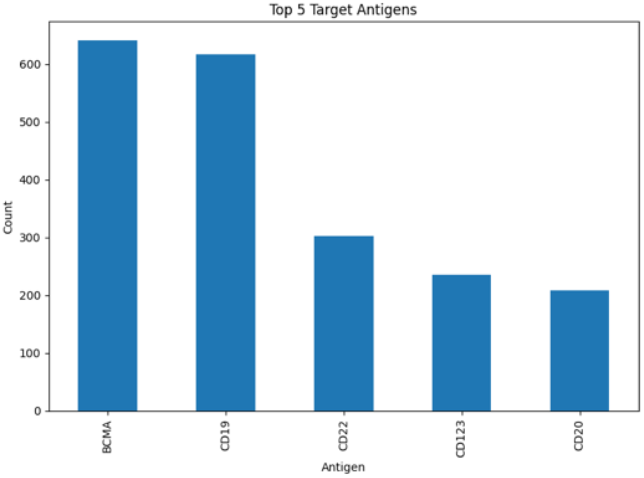
Barplot of the 5 most common target antigens in CARTiBASE.

### Diversity of transmembrane domains

To characterize the diversity of transmembrane domains used in CAR constructs, we extracted all transmembrane sequences from CARTiBASE, enumerated their frequency, and computed pairwise sequence identities across all domains.

**Table 1.**
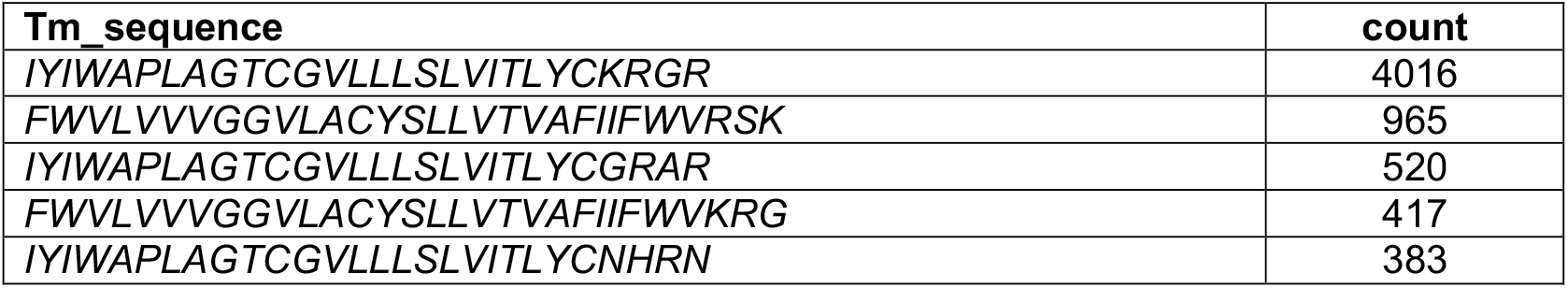
Top five transmembrane CAR sequence counts.

Hierarchical clustering reveals that the TM landscape is dominated by a small number of highly conserved sequence families (Figure 3). The most frequent TM domain, *IYIWAPLAGTCGVLLLSLVITLYCKRGR*, appears 4,016 times, far exceeding all others. Closely related variants of this TM, including *IYIWAPLAGTCGVLLLSLVITLYCGRAR* (520 occurrences) and several similar derivatives, differ by only one or two C-terminal residues, forming a tight cluster of nearly identical sequences in the heatmap. This dense cluster reflects a well-established TM backbone that has been reused across thousands of constructs.

**Figure 3.**
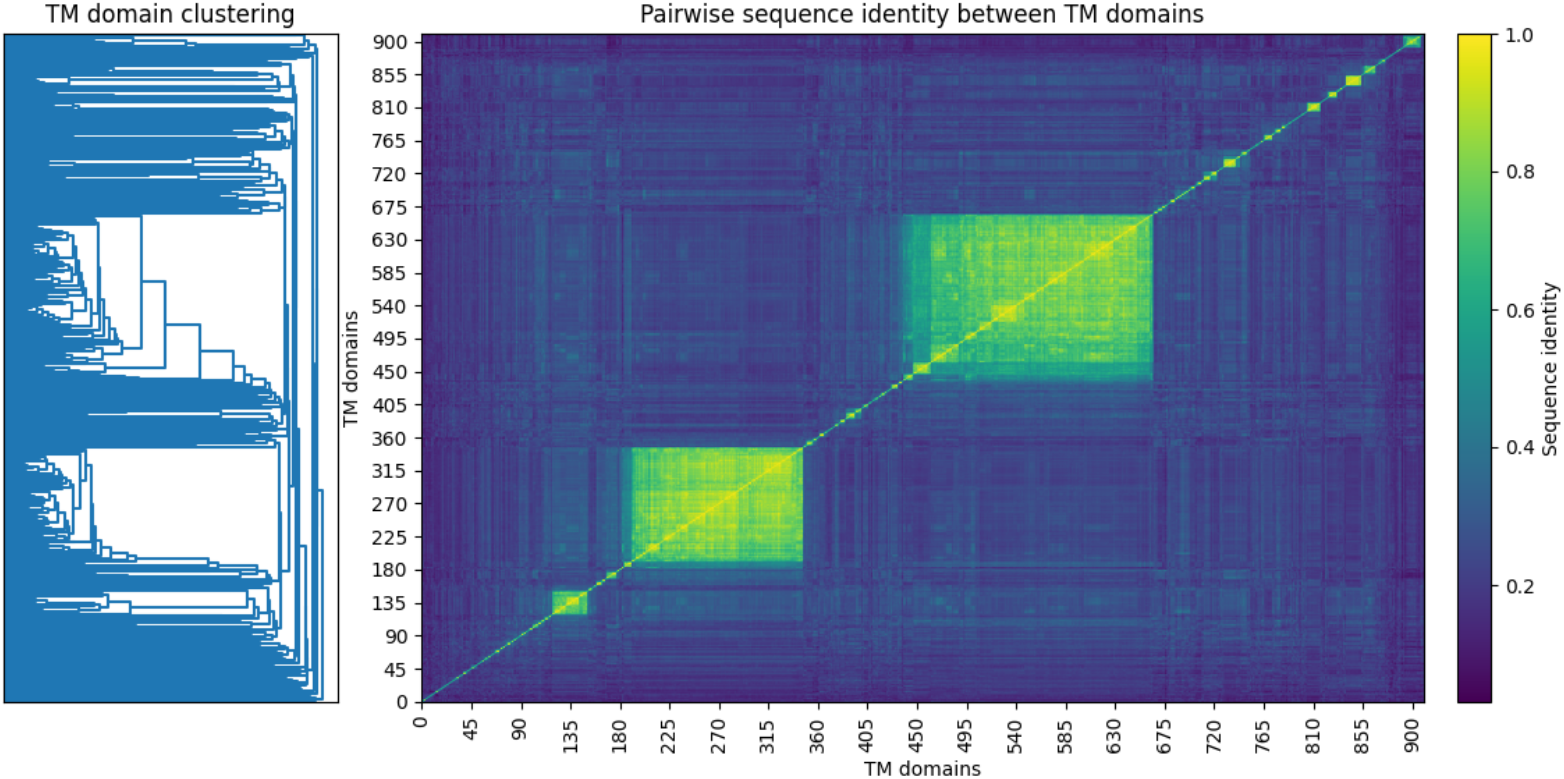
Heatmap of sequence identity between TM Domains.

A second major sequence family forms a similar block, centered around the motif *FWVLVVVGGVLACYSLLVTVAFIIFWVRS(K/R)*. These TM domains share a nearly identical hydrophobic core and vary only slightly in their charged C-terminal region, producing another high-similarity cluster that appears prominently in the matrix. Together, these two TM super-families account for the majority of all reused domains and form the two largest contiguous high-identity regions in the heatmap. The remaining TM repertoire exhibits lower overall similarity and is distributed across smaller, more heterogeneous clusters.

The clustering analysis reveals that CAR TM domains do not form a uniform or evenly sampled sequence space. Instead, they are organized into two major, tightly conserved super-families supported by extensive reuse, accompanied by a long tail of more divergent and infrequently used variants. This distribution highlights a clear design preference within the field, where a small number of canonical TM motifs serve as the backbone for most CAR constructs, while alternative TM architectures remain comparatively underexplored.

### Interpretation of BCMA scFv sequence clustering

To examine sequence diversity within the BCMA-targeting antigen-binding domains (ABDs), we computed pairwise sequence similarities for all unique scFv sequences and applied hierarchical clustering. The resulting heatmap displays the similarity matrix, with dendrograms summarizing the relationships between sequences.

Overall, the BCMA ABD repertoire forms multiple distinct sequence families. Several compact, high-similarity blocks appear along the diagonal, each representing groups of closely related scFvs that likely originate from the same parental antibody (Figure 4). These clusters correspond to well-defined lineages that have undergone only minor variation.

**Figure 4.**
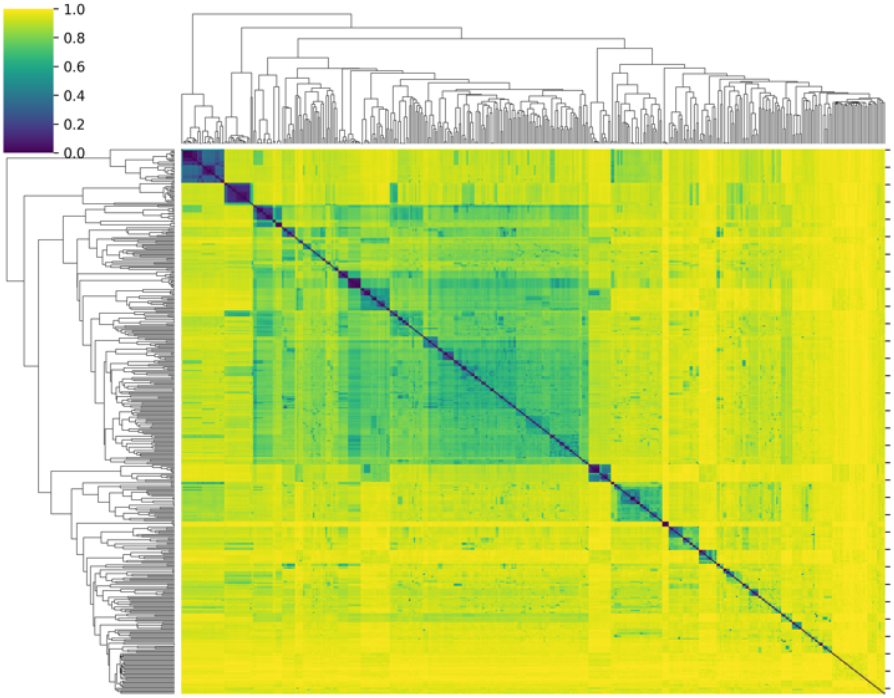
ABD similarity clustering for BCMA.

In contrast, the upper-left quadrant contains a large, diffuse green block that corresponds to a broader and more heterogeneous sequence family.

The moderate similarity values in this region indicate that these scFvs are more related to each other than to the rest of the dataset, but they are not strongly conserved. This pattern suggests a commonly used BCMA-binding scaffold that has been repeatedly modified, generating many variants through affinity maturation or CDR engineering. Rather than forming distinct subclusters, these sequences span a continuous range of variation, effectively creating a large “super-family” with considerable internal diversity. These patterns reveal that BCMA-targeting scFvs do not form a single dominant lineage but instead span multiple evolutionary or design trajectories. The presence of both tightly conserved clusters and a large heterogeneous super-family highlights the breadth of the BCMA design space.

### Interpretation of CD19 scFv sequence clustering

Within the CD19 scFv repertoire, we see multiple compact, highly similar blocks appear along the diagonal (Figure 5). These dense purple regions correspond to tight clusters of scFvs with near-identical CDR architectures, indicating the presence of several strongly conserved CD19-binding lineages. These clusters most likely represent well-established parental antibodies (e.g., FMC63-derived sequences and related variants) and their close derivatives, which dominate a portion of the CD19 design space.

**Figure 5.**
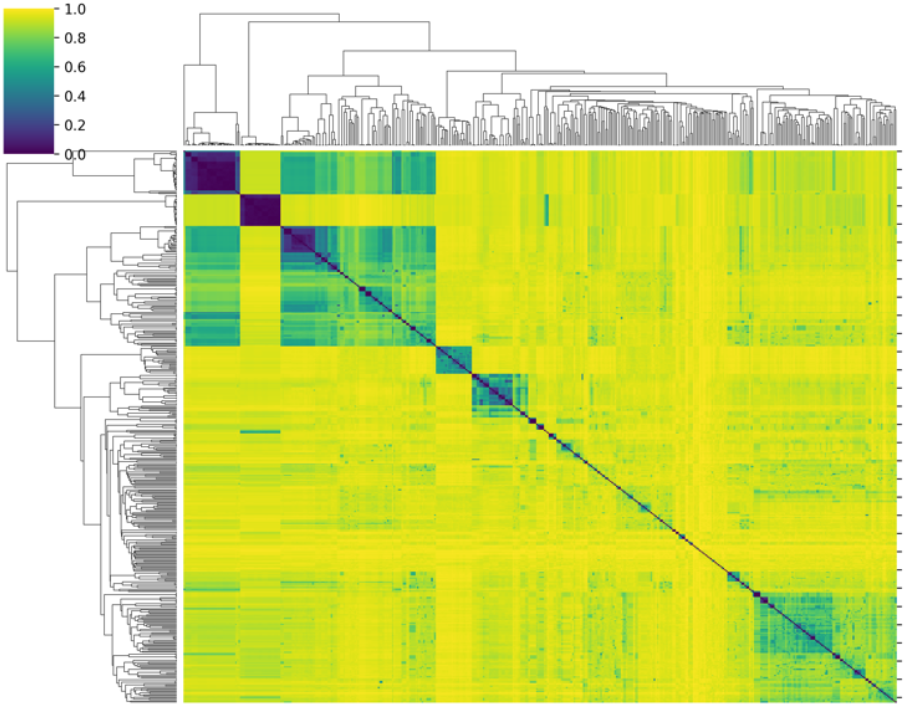
ABD similarity clustering for CD19.

Beyond these high-similarity groups, the heatmap shows a substantial diversity across the remainder of the CD19 scFv pool, with many sequences sharing only moderate similarity to one another. Consistent with this, most of the off-diagonal regions lack sharply defined blocks, suggesting that a large fraction of CD19-targeting designs arise from diversified engineering efforts.

### Interpretation of CD123 scFv sequence clustering

Analysis of the diversity within the CD123-targeting antigen-binding domains (ABDs) revealed a markedly structured landscape, dominated by a small number of highly conserved sequence lineages and a broader, heterogeneous background of more divergent variants (Figure 6). The most striking feature is the presence of two large, dark purple blocks in the upper-left quadrant. These blocks represent two highly conserved CD123-binding families. Their high intra-cluster similarity indicates that these scFvs likely derive from only a few parental antibodies that have been repeatedly reused or minimally modified across different CAR constructs. Adjacent to these conserved families, the heatmap shows a region of intermediate similarity values, indicating interactions between the major conserved clusters and a more diverse set of CD123-binding scFvs. These sequences share some degree of homology with the dominant lineages but lack the deep conservation observed within the core blocks, suggesting that they may represent engineered variants or affinity-maturation derivatives that deviate from the original scaffolds. The lower portion of the heatmap represents a large and diverse pool of CD123-targeting scFvs that do not cluster into tight families.

**Figure 6.**
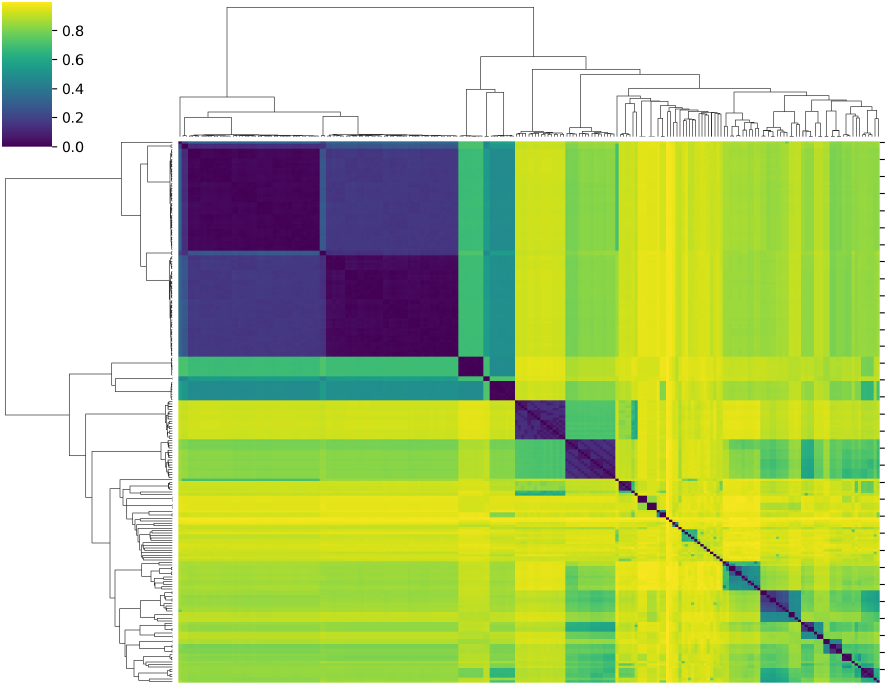
ABD similarity clustering for CD123.

## Availability and requirements

### Use case 1: retrieving reference CARs and identifying nearest neighbors

Users can retrieve CARs by providing an amino acid sequence, a patent accession number or uploading a FASTA file from which a query sequence can be selected. CARTiBASE returns the top-k most similar constructs in the database, together with their domain segmentation, associated metadata and alignment details in case the query is an amino acid sequence. This allows users to examine differences in extracellular regions, transmembrane domains and intracellular signaling modules across the closest matches, and identify common design variants such as alternative hinge lengths, transmembrane choices or different co-stimulatory architectures.

### Use case 2: exporting curated CAR data

CARTiBASE allows users to download full-length CAR sequences or domain-resolved FASTA files directly from the interface. These exports support downstream analyses such as manual inspection, custom annotation and motif discovery. The standardized domain segmentation ensures that sequences can be reused reliably across workflows without requiring additional preprocessing.

Web server: https://www.cartibase.org

Data download: FASTA exports available via the interface

License: CC BY-NC-ND

